# Integrated Metabolomics to Reveal the Impacts of Common Antibiotics based on Drug Resistance Prediction of Gut Microbiota

**DOI:** 10.1101/2021.06.16.444960

**Authors:** Pei Gao, Ming Huang, Naoaki Ono, Shigehiko Kanaya, MD Altaf-Ul-Amin

## Abstract

Due to the close interaction between the host and the gut microbiota, the alterations in gut microbiota metabolism may therefore contribute to various diseases. How to use antibiotics more wisely in clinical practice is a promising task in the field of pathophysiology related to gut microbiota. The hope fueling this research is that the alteration of gut microbial communities are paralleled by their capacity on metabolomic from the combined perspective of microbiome and metabolomics. In order to reveal the impacts of antibiotics on microbiota-associated host metabolomic phenotypes, a feasible methodology should be well developed to assess the pervasive effects of antibiotics on the population structure of gut microbial communities. Our attempt starts from predicting specific resistance phenotypes of the individuals in isolation from the rest of the gut microbiota community, according to their resistant genotypes. Once resistance phenotypes of microbiome is determined, we integrated metabolomics with machine learning by applying various analysis algorithms to explore the relationship between the predicted resistance and metabolites, including what the microbial community is after medication, which microbes produce metabolites, and how these metabolites enrich.

## Introduction

The human gut is colonized by a high abundance of microorganisms, which have been attested to be deeply integrated with human physiological function [18]. Previous researches revealed the diverse roles of gut microbiota on human physiology, including energy metabolism, behavioral and mental states, as well as pathogen defence and immune functions [8,17,19]. And products of microbial metabolism exist at a nexus between host and microbiome in a wide range of human tissues such as feces, urine, and cerebrospinal fluid [6,9]. Due to the close interaction between the host and the gut microbiota, the alterations in gut microbiota metabolism may therefore contribute to various diseases, for instance, inflammatory diseases, metabolic ailments, cancer and mental disorders.

Several host factors influence the physiological state of gut microbiota, and hence participate in the pathophysiological process of disease occurrence. Turnbaugh et al. revealed that diet-induced obesity is linked to marked but reversible alterations in the mouse distal gut microbiome [20]. Azad et al. explained hygiene hypothesis about infant allergic disease by implying the impact of hygiene on microbiota composition and diversity [3]. Besides, dozens of studies have reported that drug application exerts the protective or harmful effect against many diseases. Especially for antibiotics, increasing evidences support the correlation between their overuse and the development of many disorders associated with the alteration of gut microbiota [14]. Pinato et al. reported that broad-spectrum ATB use can cause prolonged disruption of the gut ecosystem and impair the effectiveness of the cytotoxic T-cell response against cancer [15]. Khodaie et al. found that the administration of the antibiotic minocycline can induce changes in microbiota composition and potentiate the effect of the antipsychotic drug risperidone in people with chronic schizophrenia from a double-blind study [10]. How to use antibiotics more wisely in clinical practice is a promising task in the field of pathophysiology related to gut microbiota.

Regular bench works has concerned the effects of individual antibiotics on individual, cultivated strains of bacteria in the laboratory, or on specific species of bacteria cultivated from antibiotic-exposed hosts [13]. Given that gut microbiota affect host physiology in the form of functional community, a system-wide perspective is apparently more powerful for monitoring the dynamic changes of pathological metabolic [21].

However, traditional omics analysis focus on the final variation of the disease process and intervention measures, leaving the endogenous mechanism of the change of gut microbiota and metabolite products unclear. [7]. Thus, metabolomics alone may Limits the comprehend of how antibiotics affect the metabolic state of the host by changing the gut microbiota.

From the combined perspective of microbiome and metabolomics, it can be found that the alteration of gut microbial communities are paralleled by their capacity on metabolomic [13]. In order to reveal the impacts of antibiotics on microbiota-associated host metabolomic phenotypes, a feasible methodology should be well developed to assess the pervasive effects of antibiotics on the population structure of gut microbial communities. Our attempt starts from predicting specific resistance phenotypes of the individuals in isolation from the rest of the gut microbiota community, according to their resistant genotypes. Once resistance phenotypes of microbiome is determined, we integrated metabolomics with machine learning by applying various analysis algorithms to explore the relationship between the predicted resistance and metabolites, including what the microbial community is after medication, which microbes produce metabolites, and how these metabolites enrich. And the workflow is shown in Fig. 1.

**Figure 1.**
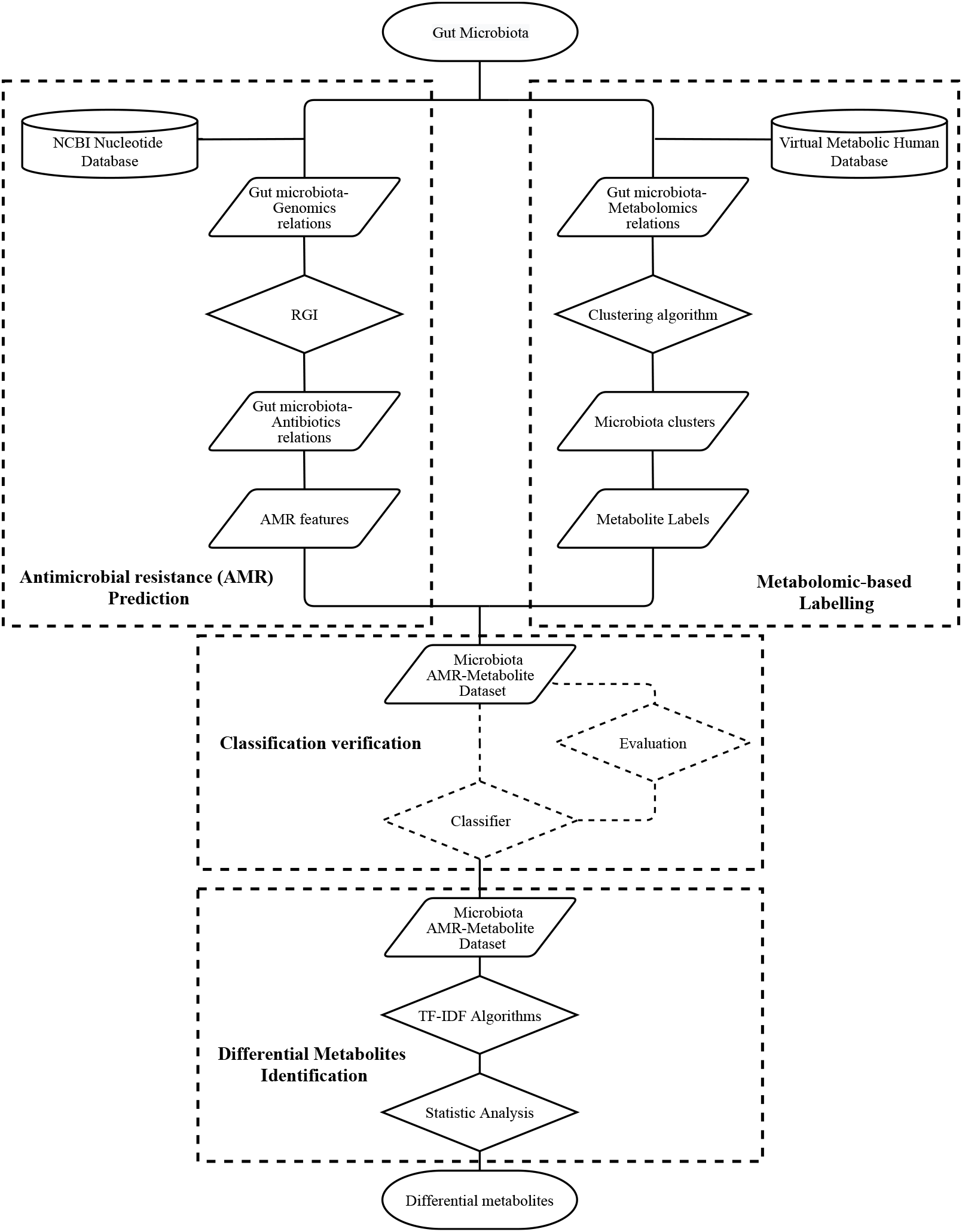
The workflow of integrated metabolomics analysis to reveal the impacts of common antibiotics based on AMR prediction of gut microbiota.

## Materials and Methods

### Data collection

We used two public databases: the Virtual Metabolic Human Database for the gut microbiota-metabolomics relations and the NCBI nucleotide database for the gut microbiota-genomics relations.

The gut microbiota resource from Virtual Metabolic Human Database contains the collection of over 800 semi-automatically curated strain-specific microbial reconstructions, belonging to 205 genera and 605 species. All microbial reconstructions were based on literature-derived experimental data and/or comparative genomics. And a typical reconstruction contains an average of 771 (±262) genes, 1198 (±241) reactions, and 933 (±139) metabolites [11]. After preprocessing, a numeric vectors of 818 bacteria and 753 metabolites were constructed to express the binary status of specific metabolite, 1 (produce) and 0 (not produce), for individual gut microbe.

The genomes & maps resource from NCBI Nucleotide Database is a collection of sequences from several sources, including GenBank, RefSeq, TPA and PDB. Genome, gene and transcript sequence data. These sequences are obtained primarily through submissions from individual laboratories and batch submissions from large-scale sequencing projects, including whole-genome shotgun (WGS) and environmental sampling projects. [4]. The sequences of 787 gut microbiota involved, mainly WGS projects, are acquired manually and formed for next step drug resistance prediction.

### Antimicrobial resistance prediction

Clinical microbiology laboratories currently perform antimicrobial susceptibility testing (AST) using phenotypic methods, such as the minimum inhibitory concentration (MIC). However, these methods are unpractical in this case because a part of gut microbiota is difficult to isolate or slow-growing for cultured. To remove the obstacle of traditional phenotypic AST based on strain-culturing, a number of molecular-level supplementary techniques have been developed. WGS provides a comprehensive inventory of an organism’s functional potential, making it an attractive approach for AST [12]. And various feasible tools and algorithms for detecting antimicrobial resistance (AMR) from WGS data are publicly accessible.

We performed AMR detection from WGS data using Resistance Gene Identifier (RGI) website portal version (RGI 5.2.0) from Comprehensive Antibiotic Resistance Database (CARD 3.1.2) [1]. CARD stands out for its high quality, manually curated resistance detection models derived from experimentally verified phenotype-genotype associations reported in the scientific literature, and through collaborations with public health and clinical microbiology laboratories [5]. When WGS of each bacteria from gut microbiota are submitted, RGI first predicts complete open reading frames (ORFs) using Prodigal and analyzes the predicted protein sequences. Resistomes from protein or nucleotide are predicted based on homology using double index alignment of next-generation sequencing data (DIAMOND), and strict significance are determined based on CARD curated bitscore cut-offs. The RGI analyzes genome or proteome sequences is under the Loose paradigms and parameters included for percent identity filtering is 50%.

### Machine Learing Algorithms

Proteomics methods that directly detect the presence and abundance of the proteins that confer AMR should, in theory, provide the strongest molecular evidence of resistance [2]. Transcriptomics approaches to AMR detection are problematic due to their low correlation with protein abundance levels. The suitability of the application of CARD for AMR prediction is promising but still questionable, and not to mention the further integration of metabolomics. For this sake, we deploy some machine learning algorithms to eliminate this uncertainty. Based on the clustering result of metabolites, we reconstructed a gut microbiota dataset with the label referring to the clustering result. And then random forest (RF) performed the contrastive classification task on predicted AMR features to evaluate the labeling. If the metabolite labeling is well-perform considering to the AMR features, it can be infer that there are some coherence between the predicted AMR and metabolite products. So that we could exert the intgrated research of gut microbiota and metabolomic to reveal the impact of antibiotics.

#### K-means

K-means is a widely used unsupervised learning algorithms for clustering. The procedure firstly set through a certain number (k) of clusters fixed a priori. The center of each cluster are defined as much as possible far away from each other. Subsequently, K centroids is recalculated until the centers are the same as the cluster centers obtained in the previous iteration. As a result of this iteration it may notice that the k centroids moves by each loop until their location are fixed.

The number of centers should be set up properly to achieve a valid clustering task on the dataset. In this case we aims at minimizing two objective function, squared error function and silhouette coefficient function.

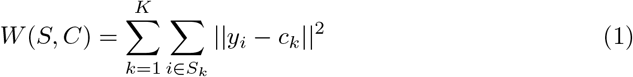

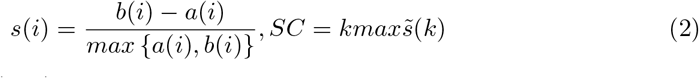

#### Random Forest (RF)

The RF is an ensemble model that grows a number of tree-based weak classifiers by randomly selecting partial bootstrapped variables. The salient features are voted by bootstrapped training process of classification task. Although each sample labeled as a binary class, the interleaving herbs that were contained in samples and the sizable feature vector will lead to an excessive variance. The RF reduces the predictive variance by decorrelating the individual weak classifier as the following equation:

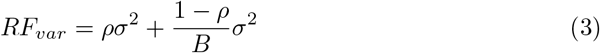

where *ρ* is the correlation of trees and *σ*^2^ is the variance of the herbs. By increasing the tree numbers *B* (the maximum is 100), the second term on the right-hand side becomes minor. Moreover, the hyperparameter max depth of a tree set as 13 in this study to avoid over fitting problem. ana

### TF-IDF Algorithms

Term Frequency Inverse Document Frequency (TF-IDF) is a commonly used weighting technique for information retrieval and data mining. The main idea of TF-IDF is to calculate values for each word in a document through an inverse proportion of the frequency of the word in a particular document to the percentage of documents the word appears in [16]. In this work, term frequency, tf(t,d), is the frequency of term t,

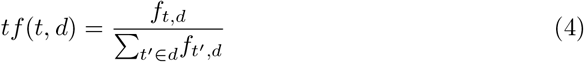

where *f*_*t,d*_ is the raw count t of bacteria containing a certain metabolite in the bacterial population d that survived the application of a certain antibiotic. And the inverse document frequency is a measure of how much information the word provides,

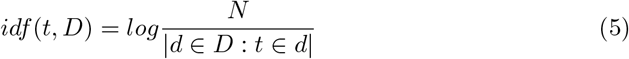

It is obtained by dividing the total number of 787 gut microbes by the number of bacteria containing the certain metabolite, and then taking the logarithm of that quotient. This algorithm aims to reflect the importance of a certain metabolite to the survival of the gut microbes after the application of a certain antibiotic.

### Metabolomics profiling

Metabolic pathway analysis was performed by MetaboAnalyst 5.0 (https://www.metaboanalyst.ca/). Pathway enrichment analysis usually refers to quantitative enrichment analysis directly based on the compound concentration values as compared to the compound lists used by over representation analysis. The available algorithms include Fishers’ exact test, the hypergeometric test, global test and Global Ancova [22]. According to the enrichment analysis, the pathways significantly affected by all the 41 antibiotics in gut microbes were displayed.

## Results

### Machine Learning Model Evaluation

To eliminate the uncertain relationship between the AMR detection of gut microbes and their metabolomics, we applied the machine learning algorithms to the gut microbe dataset. And the endogenous consistency between the predicted AMR features and the metabolite product features of the same bacteria was evaluated. The results of the machine learning algorithms are shown in Fig 2.

**Figure 2.**
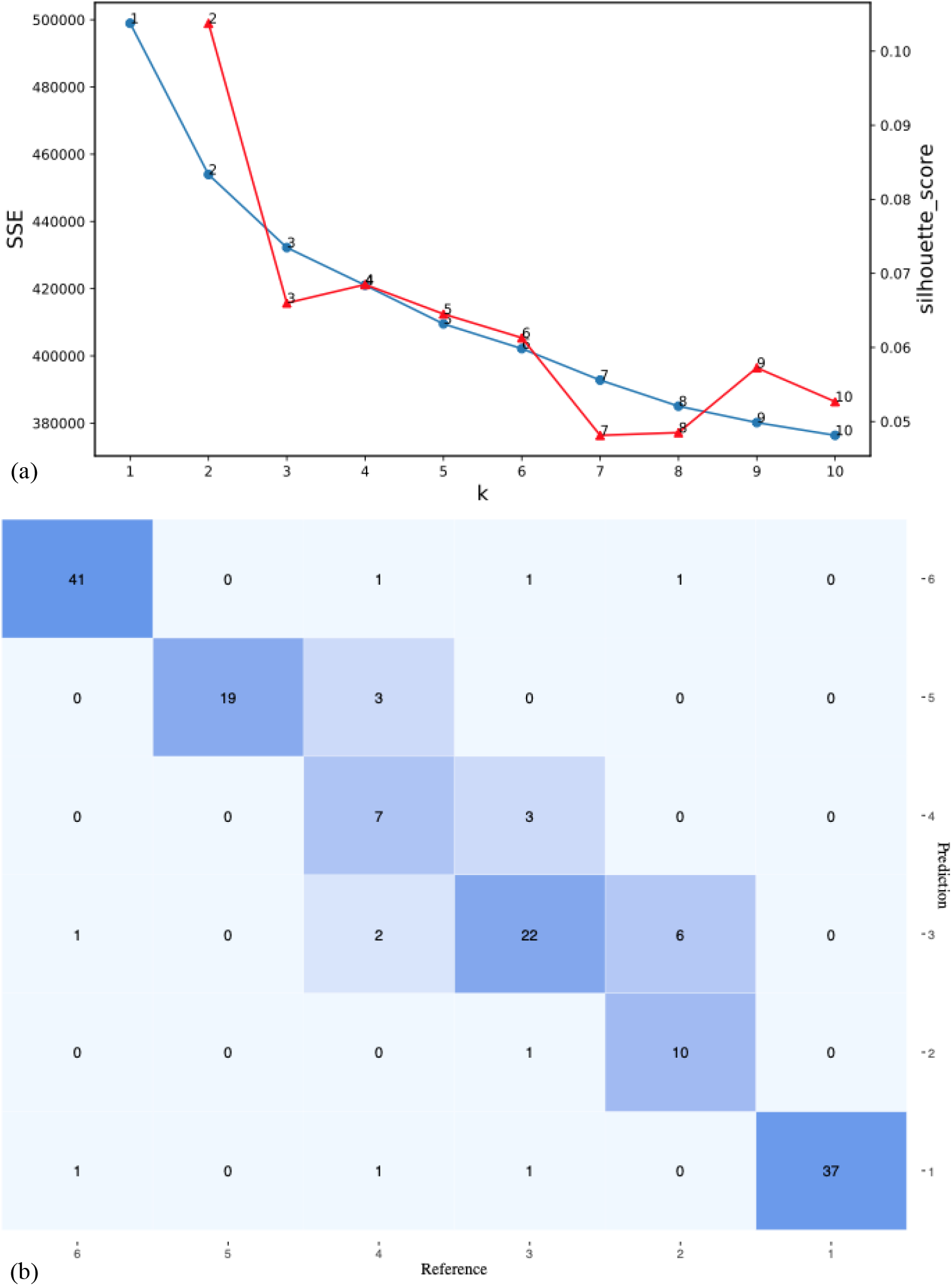
The results of machine Learning models. **(a)**, To achieve a valid clustering task on the dataset, the number of clustering centers was set up by minimizing two objective function, squared error function and silhouette coefficient function. **(b)**, Confusion matrix of RF model input with the clustering-based labelled dataset with the predicted AMR features.

Focusing on the clustering, when the number of centers set up to 6, it achieve a valid clustering task on the dataset with the metabolite product features. Two objective function, squared error and silhouette coefficient, were taken into consideration and shown in Fig 2(a). The result of the K-means was used for clustering the dataset and re-labeled all the samples, in this case, the 787 types of common gut microbes.

Focusing on the RF model, we further investigated the effect of the clustering-based label on the dataset with the predicted AMR features. The confusion matrix in Fig. 2 (b) gives more details about the RF model. It can be seen that label 2 and 6 are clearly separated without mismatching. For label 1, the subclassification is accurate, and only two misclassification occurred. Although there is a certain degree of misclassification in labels 3-5, all the specificity, according to the metrics in Table 1, are greater than 0.90. In another word, the machine learning models prove the proposed methodology for AMR prediction of gut microbiota. The dataset has latent compatibility to bridge the AMR detection of gut microbes and their metabolites. Furthermore, the selected features can be fully utilized for the next step of feature integrated metabolomics analysis.

**Table 1.**
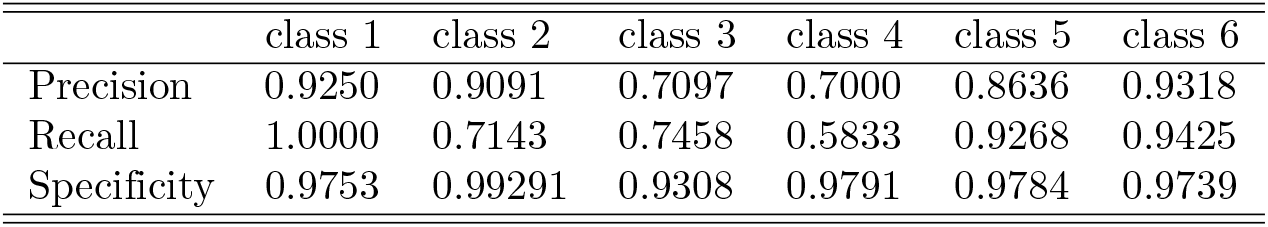
The metrics of the RF model on the clustering-based label on the dataset with the predicted AMR features.

### Differential metabolite identification

Different antibiotic classes provide different patterns of microbiota alteration because of their different spectrum and bacterial target. After the endogenous consistency between the predicted AMR and the metabolite is verified by the machine learning algorithm, the predicted AMR is methodologically proven effective. We logically mapped the resistance of the included 41 antibiotics, against a total of 787 gut microbes, into an antibacterial spectrum, as shown in Fig 3.

**Figure 3.**
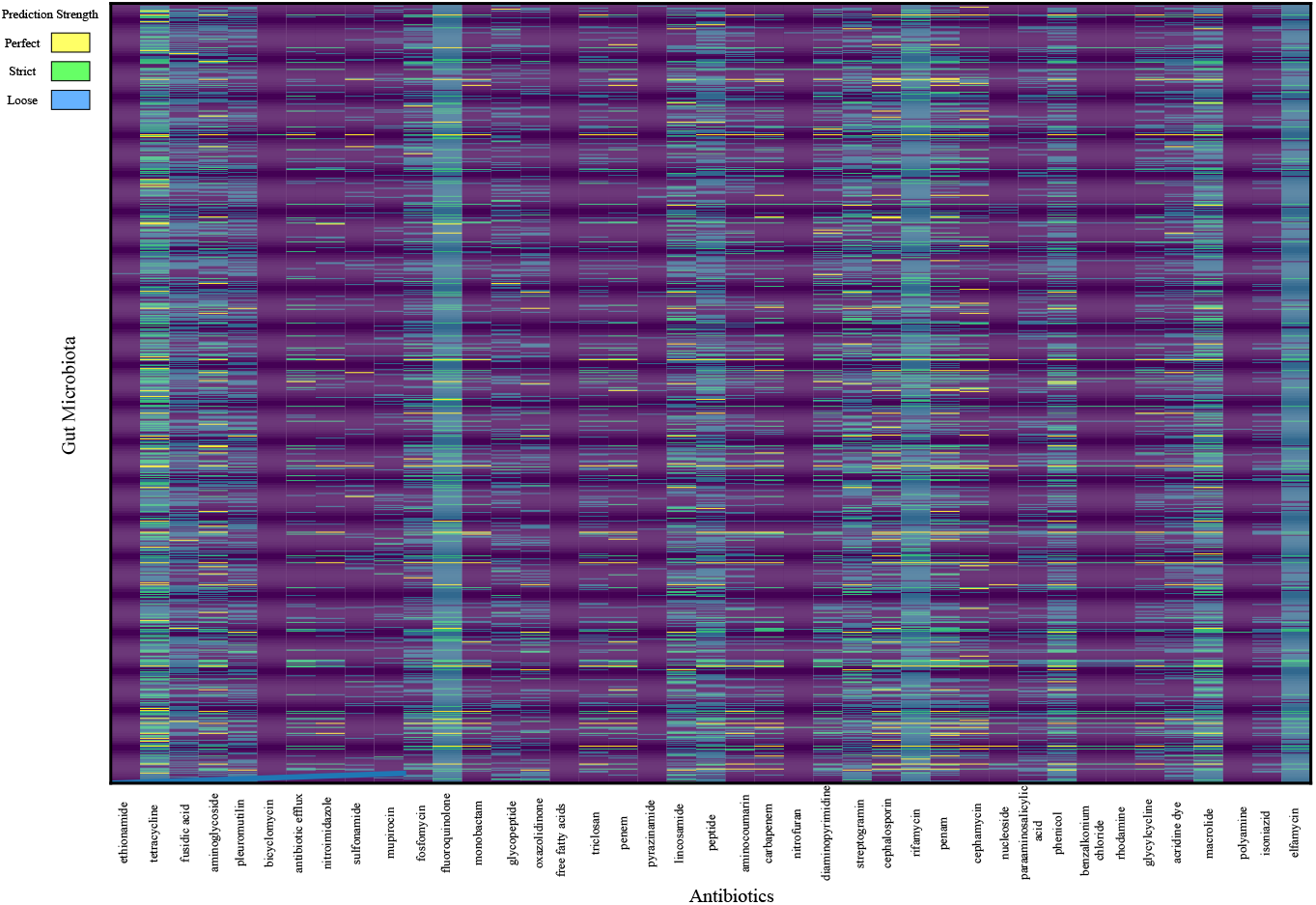
Antibacterial spectrum according to the predicted AMR. The resistance of the included 41 antibiotics, against a total of 787 gut microbes, are mapped into an antibacterial spectrum. The prediction strength of resistance to specific antibiotics are perfect (yellow), strict (green), and loose (blue).

According to the prediction strength, 6 kinds of antibiotics, ethionamide, bicyclomycin, free fatty acids, pyrazinamide, nitrofuran, and polyamine incur the least resistance, which shows consistent with the types of broad-spectrum antibiotics in clinical practice.

### Pathway enrichment analysis

To explore the affected metabolic pathways of common antibiotics in gut microbiota, we imported these differential metabolites to MetaboAnalyst 5.0. As shown in Fig 4 and Fig 5, based on pathway impact threshold which set up to 0.1, top 25 pathways affected significantly were visualized among the 41 antibiotics. Fatty acid degradation are most notably affected with 38 kinds of antibiotics, including acridine dye, aminocoumarin, aminoglycoside, antibiotic efflux, benzalkonium chloride, bicyclomycin, carbapenem, cephalosporin, cephamycin, diaminopyrimidine, elfamycin, fluoroquinolone, fosfomycin, fusidic acid, glycopeptide, glycylcycline, fusidic acid, lincosamide, macrolide, monobactam, mupirocin, nitrofuran, nitroimidazole, nucleoside, oxazolidinone, para-aminosalicylic acid, penam, penem, peptide, phenicol, pleuromutilin, polyamine, rhodamine, rifamycin, streptogramin, sulfonamide, tetracycline, and triclosan, as the related antibiotics application. For the other three kinds of antibiotics, the top affected enriched pathway are porphyrin and chlorophyll(antibacterial free fatty acids), fatty acid elongation in mitochondria (ethionamide) and butanoate metabolism (pyrazinamide).

**Figure 4.**
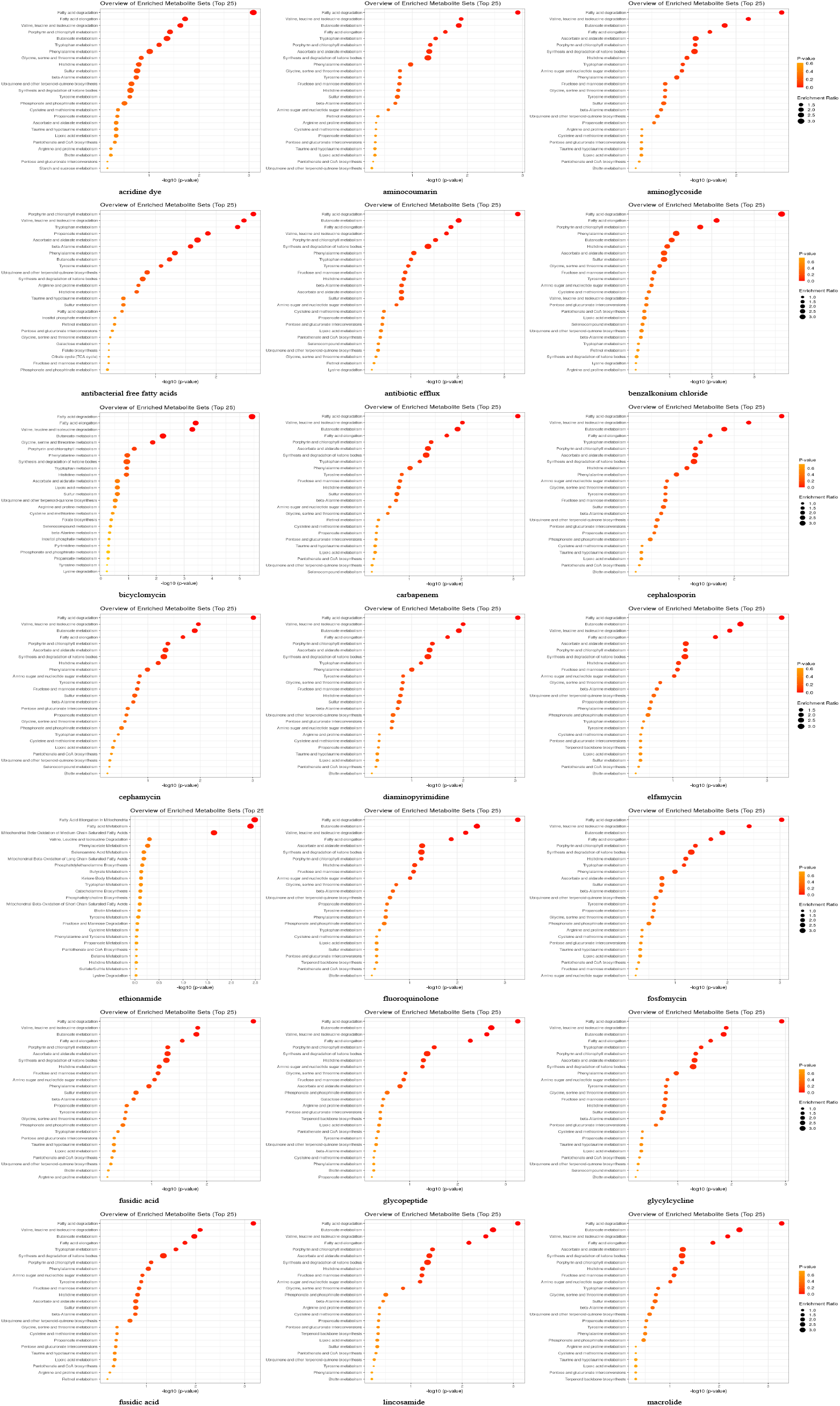
visualizing pathway enrichment analysis of the top 25 pathways affected significantly among the 41 antibiotics. The kinds of antibiotics are acridine dye, aminocoumarin, aminoglycoside, antibacterial free fatty acids, antibiotic efflux, benzalkonium chloride, bicyclomycin, carbapenem, cephalosporin, cephamycin, diaminopyrimidine, elfamycin, ethionamide, fluoroquinolone, fosfomycin, fusidicacid, glycopeptide, glycylcycline, fusidic acid, lincosamide, and macrolide.

**Figure 5.**
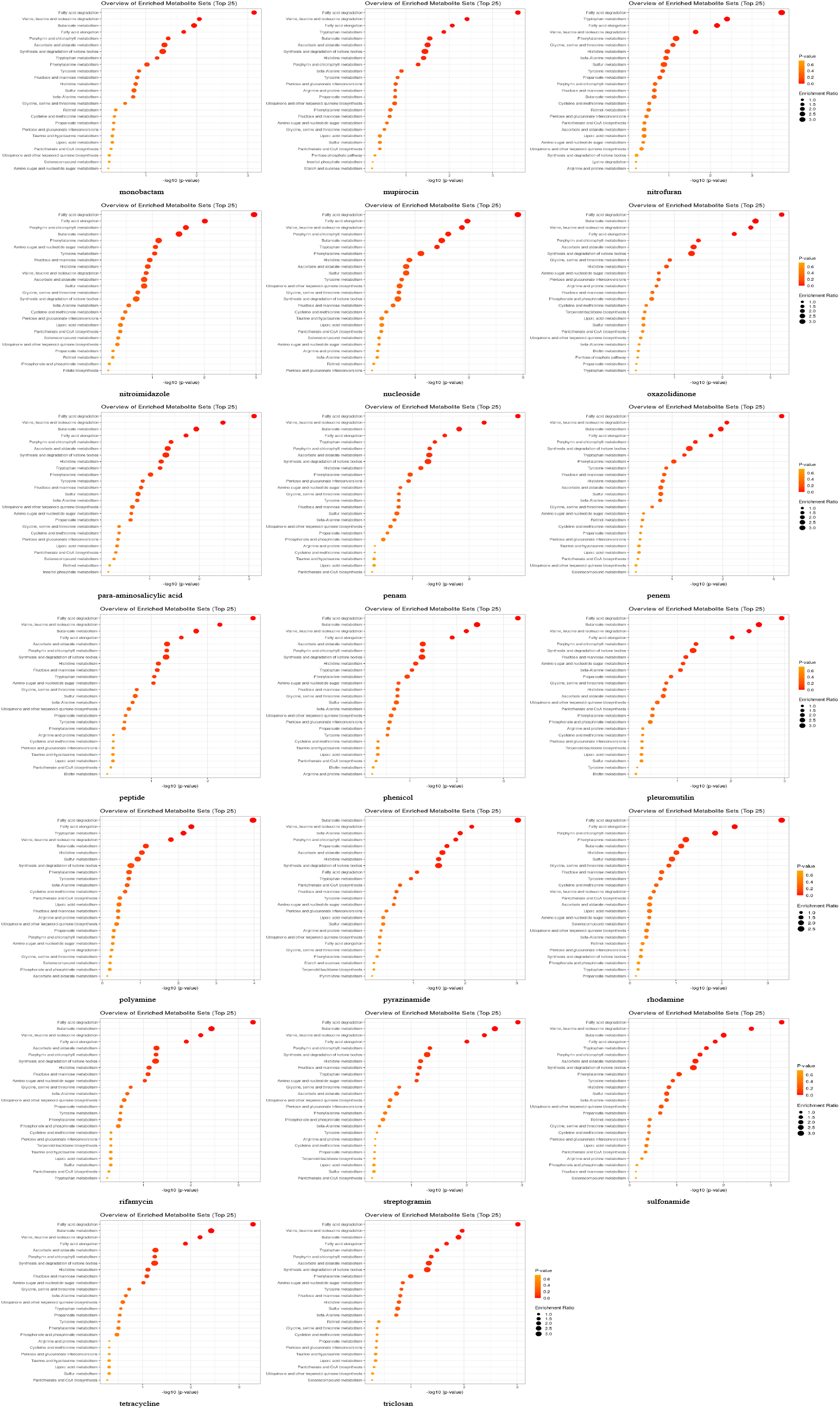
visualizing pathway enrichment analysis of the top 25 pathways affected significantly among the 41 antibiotics. The kinds of antibiotics are monobactam, mupirocin, nitrofuran, nitroimidazole, nucleoside, oxazolidinone, para-aminosalicylic acid, penam, penem, peptide, phenicol, pleuromutilin, polyamine, pyrazinamide, rhodamine, rifamycin, streptogramin, sulfonamide, tetracycline, and triclosan.

## Discussion

Commonly, a large number of bench work are indispensable when whether the gut microbiota is a valid mechanism target for disease occurrence is discussed. And whether intervention trials, in this case, administration of antibiotics, are of clinical relevance need to be examined with longitudinal trials. Nevertheless, the workflow adopted in this work can be considered as a top down approach. Because we started with the antibiotics-resistant prediction of large-scale microbiota in silico. Then we moved down to the metabolite level and utilized state of the art machine learning techniques to identify the significant consistency between predicted AMR and gut microbe metabolites. Hence the approach is also a computational approach, in which our input data corresponds to the survival of the gut microbes and the identified differential metabolites against each common antibiotics. And the results we obtained are promising showing integrated pathway enrichment, providing a breakthrough point for future mechanism research.

Another thing that is interesting to discuss is the enriched pathway regarding to the differential metabolites. The enrichment analysis show more than 90% antibiotics are related to the changed on fatty acid degradation with the greatest p values. Further research is needed to consider the directly or indirectly mechanism for taking this pathway as the potential target to bridge antibiotics application and gut microbiota-associated pathophysiology.

Last but not least, what needs to be further discussed is how the process of forming the predicted antibiotics resistance, is enough with a three-level representation as in this study or needs to be divided into the subtler classes.

## Conclusion

In this study, assuming that it can be found that the alteration of gut microbial communities are paralleled by their capacity on metabolomic, we first developed a novel integrated approach to predict the specific resistance phenotypes of the individuals in isolation from the rest of the gut microbiota community, according to their resistant genotypes. We identified the relationship between the predicted resistance and metabolites, including what the microbial community is after medication, which microbes produce metabolites, and how these metabolites enrich. The consistency between the predicted AMR and metabolomics was further validated by supervised and unsupervised machine learning methods. With the TF-IDF algorithms, the importance of a certain metabolite to the survival of the gut microbes after the application of a certain antibiotic was determined. The integrated pathway metabolomic analysis revealed fatty acid degradation are most notably affected with 38 kinds of antibiotics. It also provides a novel paradigm to identify the potential mechanisms of pharmacological effects derived from the affected gut microbiota.

## Acknowledgments

We thank just about everybody.

